# Structural mechanisms of PIP_2_ activation and SEA0400 inhibition in human cardiac sodium-calcium exchanger NCX1

**DOI:** 10.1101/2024.12.05.627058

**Authors:** Jing Xue, Weizhong Zeng, Scott John, Nicole Attiq, Michela Ottolia, Youxing Jiang

**Affiliations:** Howard Hughes Medical Institute and Department of Physiology, the University of Texas Southwestern Medical Center, Dallas, Texas, USA; Department of Biophysics, the University of Texas Southwestern Medical Center, Dallas, Texas, USA; Department of Medicine (Cardiology), UCLA, Los Angeles, California, USA; Department of Anesthesiology and Perioperative Medicine, Division of Molecular Medicine, David Geffen School of Medicine, University of California Los Angeles, Los Angeles, California, USA

## Abstract

Na^+^/Ca^2+^ exchangers (NCXs) transport Ca^2+^ across the plasma membrane in exchange for Na^+^ and play a vital role in maintaining cellular Ca^2+^ homeostasis. Our previous structural study of human cardiac NCX1 (HsNCX1) reveals the overall architecture of the eukaryotic exchanger and the formation of the inactivation assembly by the intracellular regulatory domain that underlies the cytosolic Na^+^-dependent inactivation and Ca^2+^ activation of NCX1. Here we present the cryo-EM structures of HsNCX1 in complex with a physiological activator phosphatidylinositol 4,5-bisphosphate (PIP_2_), or pharmacological inhibitor SEA0400 that enhances the inactivation of the exchanger. We demonstrate that PIP_2_ binding stimulates NCX1 activity by inducing a conformational change at the interface between the TM and cytosolic domains that destabilizes the inactivation assembly. In contrast, SEA0400 binding in the TM domain of NCX1 stabilizes the exchanger in an inward-facing conformation that facilitates the formation of the inactivation assembly, thereby promoting the Na^+^-dependent inactivation of NCX1. Thus, this study reveals the structural basis of PIP_2_ activation and SEA0400 inhibition of NCX1 and provides some mechanistic understandings of cellular regulation and pharmacology of NCX family proteins.

## Introduction

Sodium-calcium exchangers (NCX) are transporters that control the flux of Ca^2+^ across the plasma membrane and play a vital role in maintaining cellular calcium homeostasis for cell signaling^1–6^. NCXs facilitate the exchange of 3 Na^+^ for 1 Ca^2+^ in an electrogenic manner, primarily responsible for extruding Ca^2+^ from the cytoplasm. However, this process can be reversed to permit Ca^2+^ entry, depending on the chemical gradients of Na^+^ and Ca^2+^, as well as the membrane potential^3,7–13^. Three NCX isoforms (NCX1-3) are present in mammals, with each isoform bearing multiple splice variants expressed in distinct tissues, thereby modulating numerous fundamental physiological events^4,14–19^. Dysfunctions of NCXs are associated with a plethora of human pathologies including cardiac hypertrophy, arrhythmia, and postischemic brain damage^3,20–22^. The cardiac variant NCX1.1 has been extensively studied, with its function playing a central role in cardiac excitation and contractile activity^23–27^.

The eukaryotic NCX consists of a transmembrane (TM) domain with 10 TM helices and a large intracellular regulatory domain between TMs 5 and 6^4,28–33^. The TM domain is responsible for the ion exchange function in NCX. It consists of two homologous halves (TMs 1-5 and TMs 6-10) with TMs 2-3 and TMs 7-8 forming the core of the TM domain and hosting the residues that coordinate the transported Na^+^ and Ca^2+34,35^. The large regulatory domain contains two calcium-binding domains (CBD1 and CBD2). Ca^2+^ binding at CBDs enhances NCX activity and rescues it from the Na^+^-dependent inactivation, a process that manifests as slow decay of the exchange current due to elevated levels of cytosolic Na^+^ ions^12,30,31,36–41^. A stretch of residues known as XIP region (eXchanger Inhibitory Peptide) at the N terminus of the regulatory domain plays a pivotal role in the Na^+^ inactivation process^42,43^.

Several other cellular cues can also modulate NCX1 activity, including phosphatidylinositol 4,5-bisphosphate (PIP_2_) in the membrane. PIP_2_ has been shown to stimulate the NCX1 activity by reducing the Na^+^-dependent inactivation and the XIP region was suggested to participate in the PIP_2_ activation^39,44–46^. In addition, several small molecule NCX inhibitors have been developed to provide valuable tools for studying the physiological and pharmacological properties of NCXs, among which compound SEA0400 is a highly potent and selective NCX1 inhibitor that promotes the Na^+^-dependent inactivation of the exchanger^20,47–52^.

We previously determined the human cardiac NCX1 structure in an inward-facing inactivated state in which XIP and the β-hairpin between TMs 1 and 2 form a TM-associated four-stranded β-hub and mediate a tight packing between the TM and cytosolic domains, resulting in the formation of a stable inactivation assembly that blocks the TM movement required for ion exchange function^32^. The study also provides a mechanistic insight into how cytosolic Ca^2+^ binding at CBD2 destabilizes the inactivation assembly and activates the exchanger^32^. To expand our understanding of NCX modulation by PIP_2_ lipid and small molecule inhibitors, we present the cryo-EM structures of human cardiac NCX1 in complex with PIP_2_ or SEA0400, revealing the structural basis underlying PIP_2_ activation and SEA0400 inhibition of NCX1.

## Results

### PIP_2_ activation of NCX1

Phosphatidylinositol 4,5-bisphosphate (PIP_2_) has been shown to modulate NCX1 activity by reducing the Na^+^-dependent inactivation of the exchanger^39,44–46^. To characterize the effect of PIP_2_ on HsNCX1, we expressed the exchanger in Xenopus laevis oocytes and recorded the outward exchanger currents using the giant patch technique in the inside-out configuration with or without applying porcine brain PIP_2_ (Fig. 1, and Methods). The recording was performed with 12 µM free cytosolic [Ca^2+^]_i_ (bath) and the outward NCX1 current was elicited by the rapid replacement of 100 mM Cs^+^ with 100 mM Na^+^ in the bath solution. In the patches without applying PIP_2_, the outward exchanger currents quickly decay and reach a steady state due to Na^+^-dependent inactivation (Fig. 1A). Introducing 10 µM PIP_2_ at the steady state progressively increases the current that plateaus at about two-fold of the peak current measured before PIP_2_ addition (Fig. 1A). After PIP_2_ washout, the outward current remains at the pre-removal level without obvious inactivation, indicating high-affinity PIP_2_ binding and its positive modulation of NCX1 by both potentiating the peak current and reducing the Na^+^-dependent inactivation. Intriguingly, the commonly used shorter chain PIP_2_ substitute (PIP_2_ diC8) does not have the equivalent activation effect on NCX1 and the exchanger remains susceptible to Na^+^-dependent inactivation when recorded in the presence of 10 µM PIP_2_ diC8 (Fig. 1B). However, PIP_2_ diC8 still binds and stimulates both the peak and steady currents of the exchanger. This stimulation effect is abolished after PIP_2_ diC8 washout, indicating a lower affinity of short-chain PIP_2_ than that of the long-chain native PIP_2_.

**Fig. 1.**
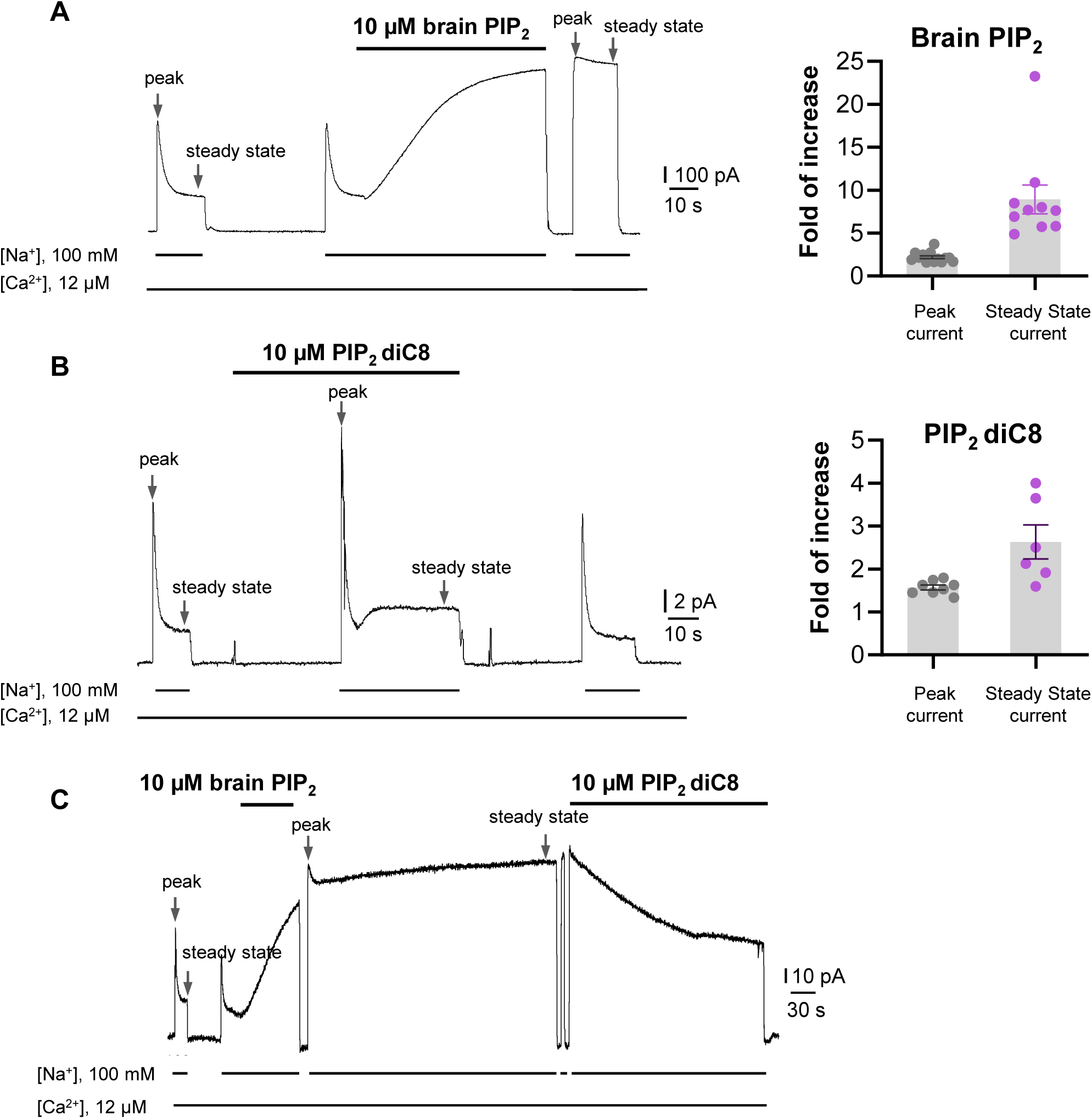
PIP_2_ enhances HsNCX1 activity. **(A)** Representative outward currents recorded from oocytes expressing the human NCX1 before and after application of long-chain brain PIP_2_. Currents were activated by replacing cytosolic Cs+ with Na+. Application of 10 µM brain PIP_2_ enhanced HsNCX1 current and abolished the Na+-dependent inactivation irreversibly. Perfusion time of PIP_2_ is indicated above traces while lines below traces indicate solution exchange. Arrows mark the peak and steady currents used to measure the fold of increase upon PIP_2_ application. The fold of current increase was calculated by comparing the peak or steady-state current before and after PIP_2_ application. **(B)** Representative outward currents recorded before and after application of short-chain PIP_2_ diC8 (10 µM). PIP_2_ diC8 was perfused from the cytosolic side before HsNCX1 activation (in the presence of Cs+ for 30 s) and during transport (in the presence of Na+). Both peak and steady-state currents of HsNCX1 are enhanced by PIP_2_ diC8 and the effect is reversible. The Na+-dependent inactivation remains in the presence of PIP_2_ diC8. **(C)** Representative outward currents recorded with the application of brain PIP_2_ and PIP_2_ diC8. The NCX1 current was first potentiated by applying 10 µM brain PIP_2_ at the steady state. The PIP_2_ effect was not reversible over the 5-minute washout with a solution containing 100 mM Na+ and 12 µM Ca2+. The same patch was then perfused with the same solution in the presence of 10 µM PIP_2_ diC8. Application of the short-chain PIP_2_ diC8 facilitates the decrease of brain PIP_2_-potentiated current, suggesting that both lipids compete for the same binding site.

To verify that the short-chain PIP_2_ diC8 and the long-chain brain PIP_2_ share the same binding site, we performed a competition assay. As shown in Fig. 1C, introducing high-affinity brain PIP_2_ at the steady state yields a long-lasting potentiation of NCX1 current that is irreversible even after a 5-minute washout. Applying PIP_2_ diC8 can steadily decrease the brain PIP_2_-potentiated NCX1 current, suggesting that both lipids compete for the same binding site.

### Structural insight into PIP_2_ binding in NCX1

To reveal the structural mechanism of PIP_2_ activation, we tried to obtain the EM structure of HsNCX1 in the presence of the long-chain porcine brain PIP_2_. However, the exchanger becomes highly dynamic, yielding a low-resolution EM map with an overall shape similar to a cytosolic Ca^2+^-activated NCX1 whose β-hub-mediated inactivation assembly is destabilized and cytosolic domain (CBD1 and CBD2) is detached from the TM domain^32^ (Fig. S1). We suspect the long-chain PIP_2_ exerts the same activation effect on NCX1 as high cytosolic Ca^2+^ by destabilizing the inactivation assembly. As NCX1 retains its Na^+^-dependent inactivation property in the presence of the short-chain PIP_2_, we reasoned that the PIP_2_ diC8-bound NCX1 likely remain in an inward-facing inactivated state in high Na^+^ low Ca^2+^ condition and its structure would still reveal how PIP_2_ binds in NCX1. We therefore determined the EM structure of NCX1 in complex with PIP_2_ diC8 at 3.5Å (Fig. 2, Fig. S2, Table S1, and Methods), which indeed adopts an inward-facing conformation with intact inactivation assembly. Due to the relative dynamic movement between the TM and cytosolic domains, we also performed local refinement to improve the map quality for each domain (Fig. S2). The density of the IP_3_ head group from the bound PIP_2_ diC8 is well-defined in the local-refined EM map focused on the TM domain (Fig. 2A and Fig. S2B). This density is not present in the apo NCX1 structure (Fig. S3). The acyl chains, however, are flexible and could not be resolved in the structure (Fig. S2). The lipid is attached to the cytosolic sides of TMs 4 and 5 with its head group positioned at the C-terminal end of TM5 (Fig. 2A). Four positively charged residues including K164 and R167 from the N-terminus of TM4 and R220 and K225 from the C-terminus of TM5 are positioned in the vicinity of the PIP_2_ head group and likely participate in the electrostatic interactions with the head group (Fig. 2A).

**Fig. 2.**
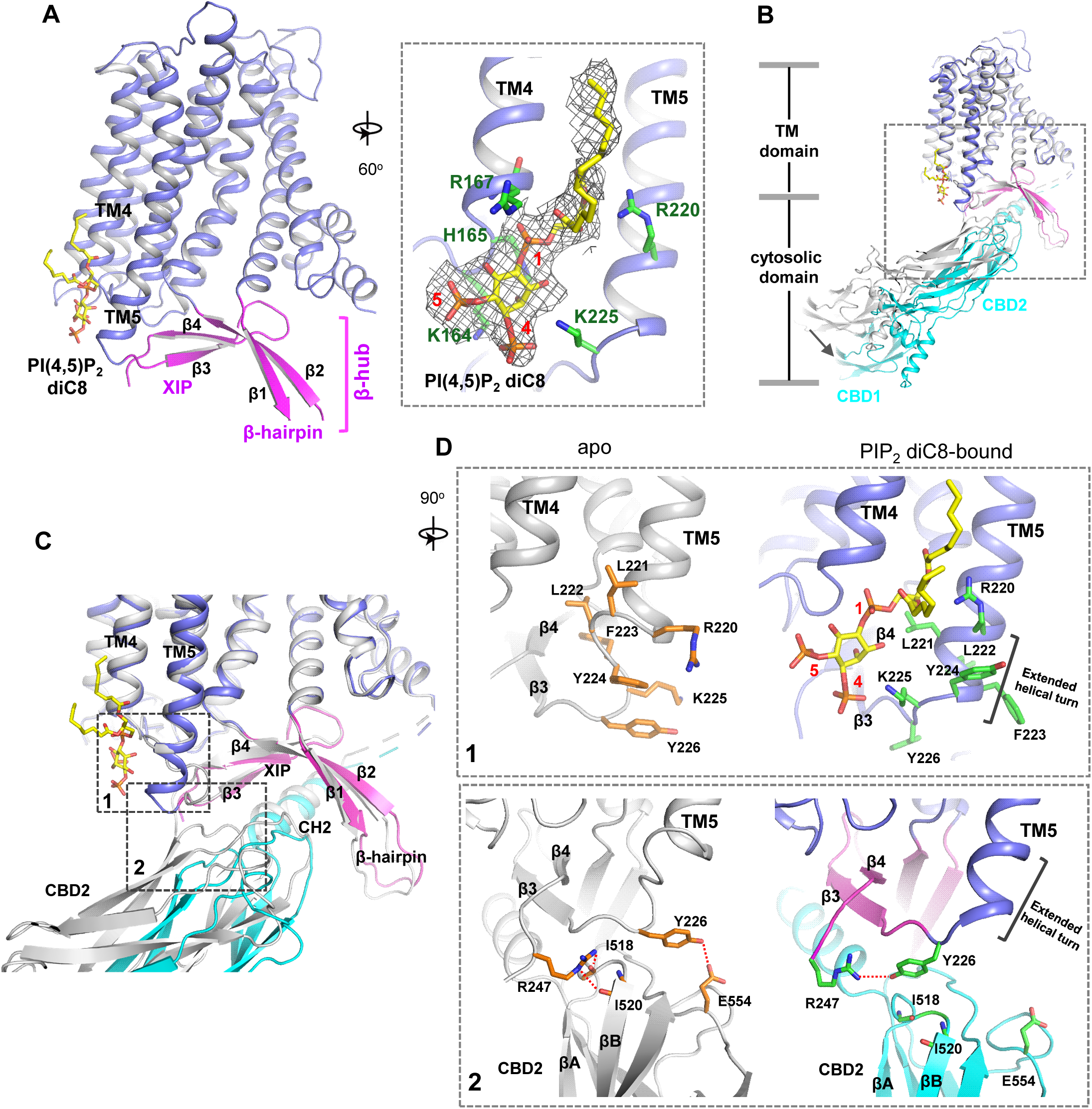
PIP_2_ binding in NCX1. **(A)** Structure of the TM and β-hub regions of PIP_2_ diC8-bound NCX1 with a zoomed-in view of the lipid binding site. The density of PIP_2_ diC8 is shown as a grey mesh contoured at 5.5*σ*. **(B)** Structural comparison between apo (grey) and PIP_2_ diC8-bound (color) NCX1. Upon PIP2 diC8 binding, there is a rigid-body downward swing movement (marked by an arrow) at CBDs caused by the partial detachment of the CBD2 domain from XIP. The conformational change at the TM domain is subtle and mainly occurs at the C-terminus of TM5 as illustrated in (**C**) and (**D**). **(C)** Zoomed-in view of the structural comparison (boxed area in (B)). The two major conformational changes occur in the boxed regions. **(D)** Zoomed-in views of the two conformational changes between apo (left in grey) and PIP_2_ diC8-bound (right in color) state. Top: conformational change 1 at the C-terminus of TM5. Bottom: conformational change 2 at the interface between XIP and CBD2.

### PIP_2_ diC8-induced conformational changes in NCX1

Two major conformational changes occur in NCX1 upon PIP_2_ diC8 binding (Fig. 2B-D). The first change occurs at the C-terminus of TM5 which ends at R220 and is connected to the 2-stranded XIP β-sheet (β3 and β4) via a 6-residue loop in the apo structure. When PIP_2_ binds, the TM5 helix extends by one helical turn. This loop-to-helix transition significantly changes the locations of these connecting loop residues and their side-chain orientations to accommodate PIP_2_, enabling K225 to reorient and interact with the IP_3_ head group (Fig. 2D, top panel). The second conformational change is the partial detachment of the CBD2 domain from XIP upon PIP_2_ binding, resulting in a downward swing of cytosolic CBD domains (CBD1 and CBD2) as a rigid body (Fig. 2B&C). This CBD2 detachment is a result of the first PIP_2_-induced conformational change at TM5 that disrupts part of the interactions between CBD2 and XIP as follows: In the apo inactivated state, Y226 and R247, the two termini residues of the 2-stranded XIP β-sheet, form H-bonds with several CBD2 residues including E554 sidechain and backbone carbonyls of I518 and I520 (Fig. 2D, bottom panel); The loop-to-helix transition of TM5 upon PIP_2_ binding leads to a dramatic rotation of Y226 that allows it to move closer to and directly interact with R247, resulting in the loss of H-bonding interactions between XIP and CBD2 (Fig. 2D, bottom panel); In addition, the rotation of Y226 also causes a direct collision with CBD2 if it remains closely attached to XIP. Thus, the PIP_2_-induced TM5 movement, particularly the reorientation of Y226, abolishes some local interactions between CBD2 and XIP and also pushes CBD2 away from XIP, causing the partial detachment of CBD2. However, the short-chain PIP_2_ only partially destabilizes rather than completely disassembles the inactivation assembly as the CH2 helix of CBD2 still engages in extensive interactions with the C-shaped β-hub.

To test if the PIP_2_-interacting residues play a critical role in the native long-chain PIP_2_ activation, we performed mutagenesis to those positively charged residues, including K164A, R167A, R220A, and K225A single mutants. We also generated an R220A/K225A double mutant as these two residues undergo PIP_2_-induced conformational change at TM5. While all mutants remain susceptible to current potentiation upon brain PIP_2_ application as seen in the wild-type NCX1, the PIP_2_-potentiation effects on peak and steady-state currents are weakened in some mutants, most notably in R220A and R220A/K225A (Fig. 3A-C). Interestingly, these two mutants also have reduced Na^+^-dependent inactivation as demonstrated by their higher fractional activity (ratio between stead-state and peak currents) before applying PIP_2_ (Fig. 3D). The retained PIP_2_ activation on these mutants implies that the longer acyl chain from native PIP_2_ plays an important role in its interaction and activation of NCX1. Furthermore, as the interactions between PIP_2_ and NCX1 are both electrostatic involving multiple charged residues and hydrophobic involving the long lipid acyl chain, those single or double amino acid substitutions may only decrease the affinity of PIP_2_ rather than abolish its binding. To test that, we also mutated all four positively charged residues to alanine. The K164A/R167A/R220A/K225A mutant is no longer sensitive to PIP_2_ activation. The currents from this quadruple mutant are small in most recordings and show no Na^+^-dependent inactivation. The unresponsiveness to PIP_2_ and lack of Na^+^-dependent inactivation in this mutant is consistent with previous studies demonstrating that PIP_2_ activates NCX by tuning the amount of Na^+^-dependent inactivation and any mutation that decreases NCX sensitivity to PIP_2_ will also affect the extent of Na^+^-dependent inactivation^44^. This quadruple NCX1 mutant likely abolishes the potentiation effect from both endogenous and externally applied PIP_2_ and thereby functions in a Na^+^-inactivated steady state.

**Fig. 3.**
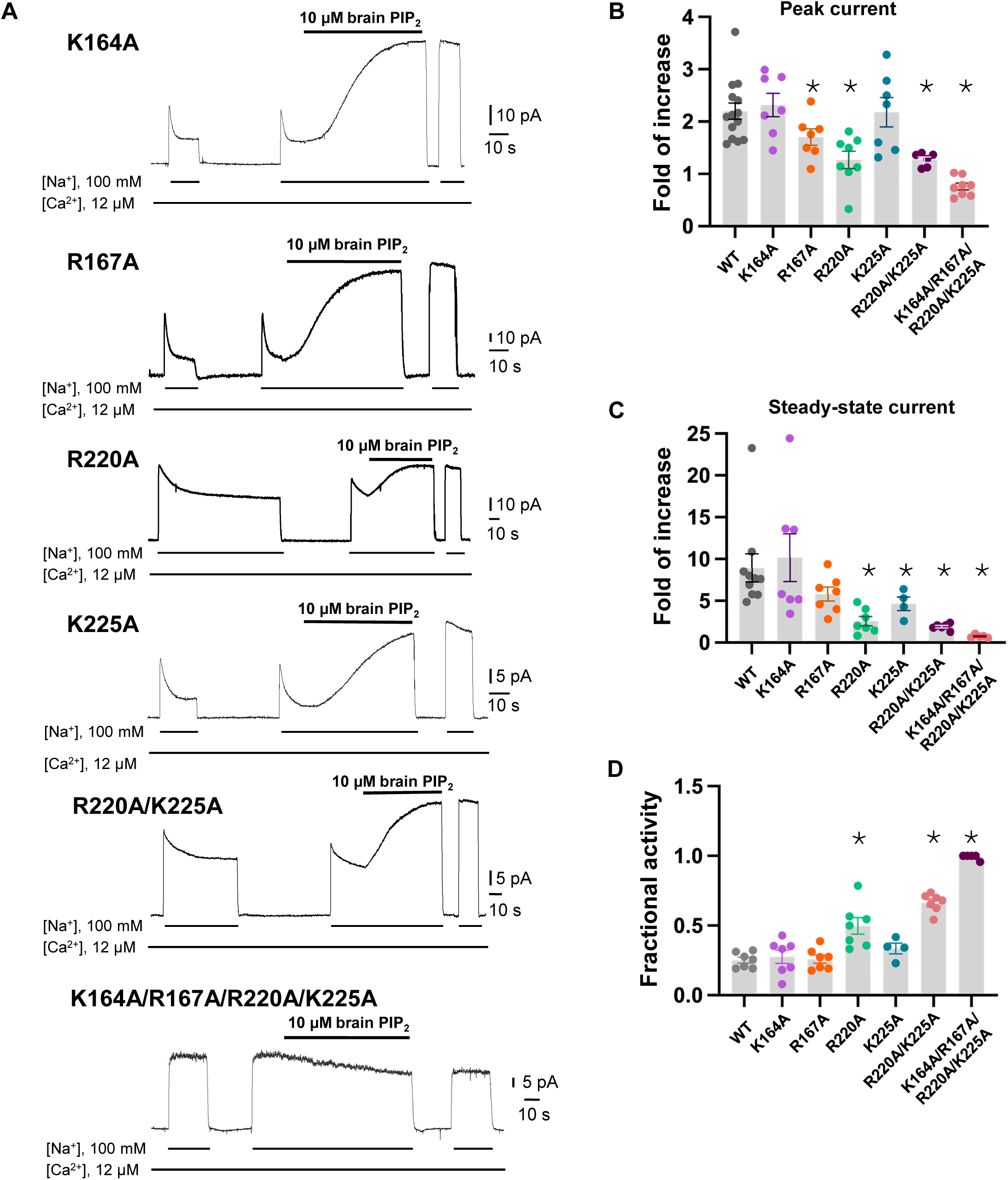
Mutagenesis at the PIP_2_ binding. **(A)** Representative NCX1 currents of PIP_2_ site mutants before and after perfusion of 10 µM brain PIP_2_ to the cytosolic side of the patch. **(B&C)** Summary graphs demonstrating the effects of PIP_2_ on the enhancement of peak (B) and steady-state currents (C). Potentiation (fold of increase) was measured by comparing the current magnitude before and after PIP_2_ application. Mutants R167A, R220A, and K225A showed some decreased response to PIP_2_ whereas R220A/K225A mutant shows more profound decrease in PIP_2_ response. Compared to WT, the PIP_2_ potentiation of R220A/K225A at the steady state is decreased by ∼70-90% (Fold of increase WT=8.9±1.6, n=10 vs R220A/K225A=1.9±0.1, n=6). PIP_2_ has no potentiation effect on the quadruple K164A/R167A/R220A/K225A mutant. Data points are mean ± s.e.m. (* p <0.1). **(D)** The extent of Na+-dependent inactivation was measured as the ratio between steady state and peak currents (fractional activity) and values for WT and the indicated mutants are shown. Mutants R220A and R220A/K225A displayed significantly higher fractional activity values when compared to WT, indicating that the Na+-dependent inactivation was less pronounced in these mutant exchangers. K164A/R167A/R220A/K225A shown no Na+-dependent inactivation.

### Structure of NCX1 in complex with SEA0400 inhibitor

SEA0400 is known to potently inhibit cardiac NCX1 by facilitating the inactivation of the exchanger^50–52^. To reveal the structural mechanism of its inhibition, we determined the structure of HsNCX1 in complex with SEA0400 at 2.9 Å (Fig. 4, Fig. S4, Table S1, and Methods). The SEA0400-bound NCX1 structure adopts an inward-facing, inactivated state identical to the apo NCX1 structure obtained at high Na^+^, nominal Ca^2+^-free condition ^32^ (Fig. 4A and Fig. S5). SEA0400 binds at the TM domain in a pocket enclosed by the internal halves of TMs 8 (8a segment), 2 (2ab segments), 4, and 5 (Fig. 4B). The pocket has a lateral fenestration in the middle of the membrane (Fig. 4B), which provides a portal for the SEA0400 entrance. The pocket is sealed off at the cytosolic side by E244 from the XIP β-sheet. As demonstrated in our previous study^32^, XIP and the linker β-hairpin (β1 and β2) between TMs 1 and 2ab form a β-hub that stabilizes the exchanger in an inactivated state. This β-hub has to be dissembled in an activated NCX1 and XIP is expected to be detached from the TM domain, which would lead to the opening of the SEA400-binding pocket to the cytosol and provide a cytosolic portal for the release of the inhibitor release as further discussed below. Fig. 4C summarizes the interactions between SEA0400 and NCX1 and demonstrates that SEA0400 fits perfectly in the pocket, making extensive contact with surrounding residues. Mutations of some key interacting residues, such as F213 and G833, have been shown to compromise the inhibitor binding^52^. The same SEA0400 binding was also demonstrated in the recent study of human NCX1.3 and mutations at some pocket-forming residues mitigate the SEA400 inhibition of the exchanger^33^.

**Fig. 4.**
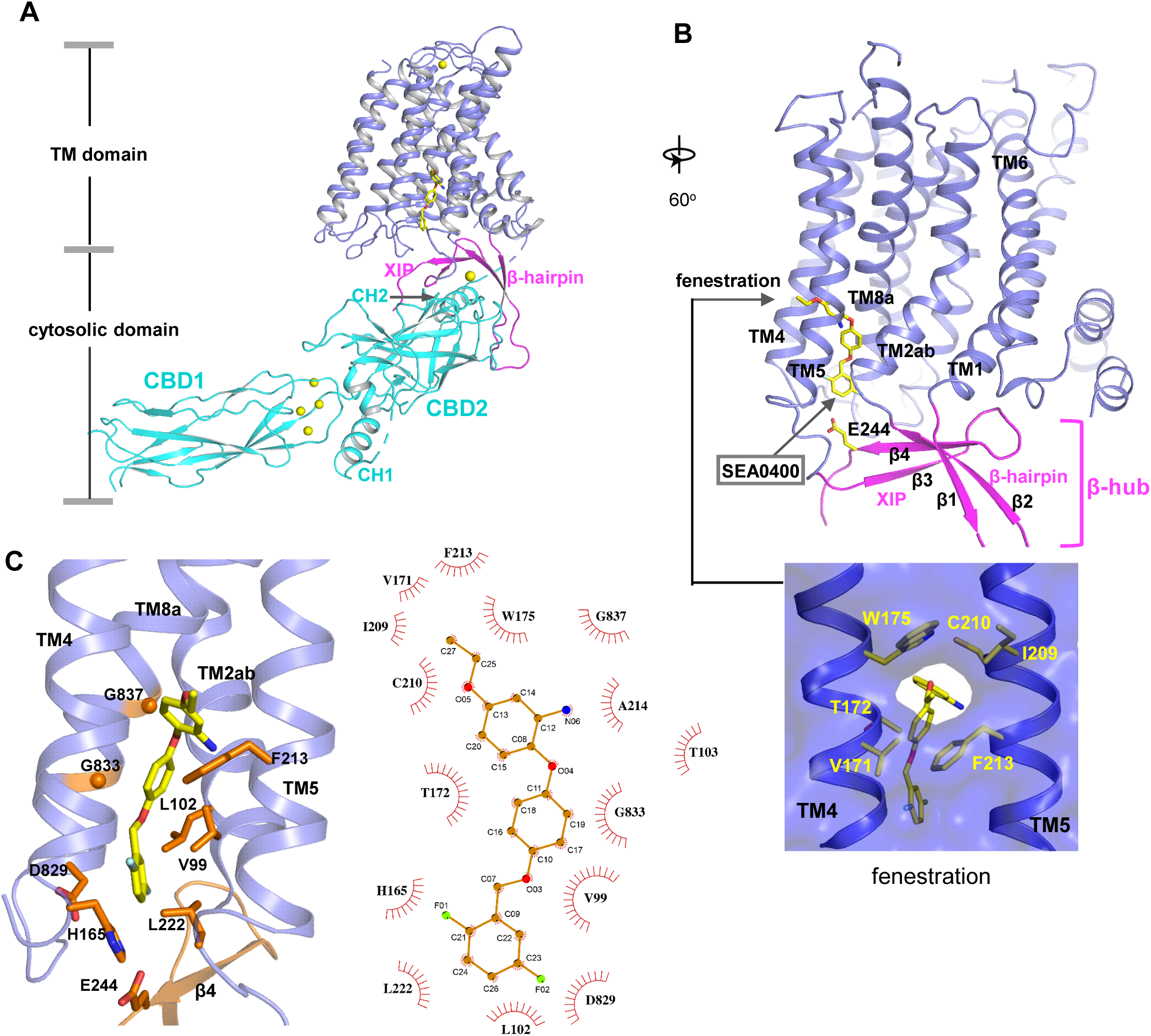
SEA0400 binding in NCX1. **(A)** Overall structure of human NCX1 in complex with SEA0400 obtained in high Na+ and low Ca2+ conditions. Yellow spheres represent the bound Ca2+ in CBD1 and XIP. **(B)** Cartoon representation of the TM domain and β-hub of the complex with surface-rendered view of the fenestration in the middle of the membrane. The β-hub is assembled by β-hairpin (β1 and β2) and XIP (β3 and β4). **(C)** Zoomed-in view of SEA0400 binding site and the schematic diagram detailing the interactions between NCX1 residues and SEA0400.

### Inhibition mechanism of SEA0400

The TM domain of NCX1 shares a similar overall architecture to the archaea exchanger NCX_Mj^34^. The structural comparison between the TM domains of the inward-facing NCX1 and the outward-facing NCX_Mj reveals the conformational changes that occur during ion exchange, providing structural insight into the SEA0400 inhibition mechanism (Fig. 5A). The inward-outward transition mainly involves the sliding motion of TMs 1 and 6 and the bending movement of TMs 2ab and 7ab^32,34,35,53^. As TM2ab directly interacts with SEA0400 in the inward-facing state, its bending movement towards the outward conformation would cause a direct collision with the inhibitor (Fig. 5A). Thus, the binding of SEA0400 stabilizes the exchanger in the inward-facing state and blocks the conformational change from the inward to the outward state. The Na^+^-dependent NCX1 inactivation occurs when the exchanger is in a Na^+^-loaded, inward-facing state with low cytosolic [Ca^2+^]^40,41,54^. As demonstrated in our previous study, only in this state, the β-hub can form and readily interact with the cytosolic CBD domains, generating the inactivation assembly that locks TMs 1 and 6 and prevents the TM module from transporting ions^32^. Thus, SEA0400 promotes NCX1 inactivation by stabilizing NCX1 in the inward-facing conformation which facilitates the formation of the inactivation assembly. The formation of the inactivation assembly also reciprocally stabilizes SEA0400 binding as the XIP of the assembly interacts with the TM domain and seals off the inhibitor binding pocket from the cytosolic side (Fig. 5B). Under conditions in which the inactivation assembly cannot form in NCX1, such as chymotrypsin treatment or forward exchange mode (Na^+^ influx/Ca^2+^ efflux), the removal or detachment of XIP from the TM domain would generate a cytosolic open portal for SEA0400 release and effectively reduce its binding affinity (Fig. 5C)^50,51^. Indeed, cysteine scanning mutagenesis studies have shown that the pocket-forming G833 residue is accessible to intracellular sulfhydryl reagents in the chymotrypsin-treated inward-facing NCX1, confirming the cytosolic exposure of the SEA400-binding pocket upon XIP removal^55^. This cytosolic opening of the SEA0400 pocket explains the low efficacy of SEA0400 inhibition in NCX1 when the Na^+^-dependent inactivation is absent.

**Fig. 5.**
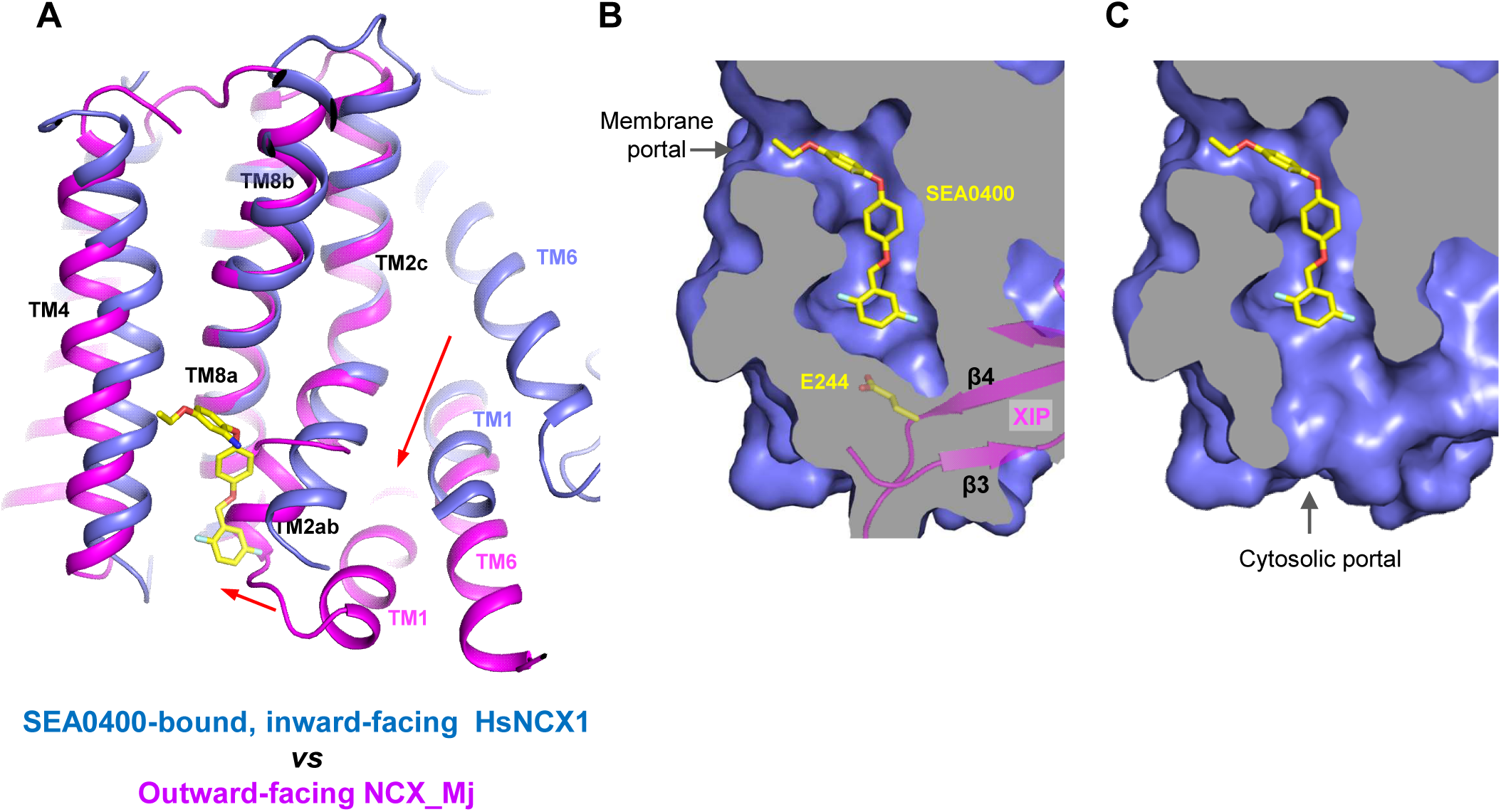
Structural mechanism of SEA0400 inhibition. **(A)** Structural comparison at the core part of the TM domain between the SEA0400-bound, inward-facing HsNCX1 and the outward-facing NCX_Mj (PDB 3V5U). Red arrows mark the sliding movement of TMs 1 and 6 and the bending of TM2ab from inward to outward conformation. **(B)** Surface-rendered views of the SEA0400-binding pocket sealed off from the cytosolic side by E244 from XIP in the inactivated state. **(C)** Removal of XIP would generate a cytosolic portal that facilitates the release of SEA0400.

## Discussion

In this study, we provide mechanistic underpinnings of cellular regulation and pharmacology of NCX family proteins by revealing the structural basis of PIP_2_ activation and SEA0400 inhibition in HsNCX1. Both compounds modulate NCX1 activity by reducing or enhancing the physiologically relevant Na^+^-dependent inactivation process that occurs when the exchanger is in an inward-facing, Na^+^-loaded state with high [Na^+^] and low [Ca^2+^] on the cytosolic side^27,40^. In this inactivation state, XIP and β-hairpin can assemble into a TM-associated four-stranded β-hub that mediates a tight packing between the TM and cytosolic domains, resulting in the formation of an inactivation assembly that blocks the TM conformational changes required for ion exchange function^32,33^. PIP_2_ or SEA0400 binding changes the stability of the inactivation assembly in NCX1, resulting in a reduction or potentiation of inactivation.

Although short-chain PIP_2_ cannot fully recapitulate the NCX1 activation effect from long-chain native PIP_2_, the structure of PIP_2_ diC8 NCX1 allows us to define the lipid-binding site and the lipid-induced conformational changes at the interface between CBD2 and XIP that destabilize the inactivation assembly. The SEA0400-bound NCX1 structure presented here, along with the recent study by Dong et al.^33^, suggests that the drug can directly inhibit NCX1 by blocking the inward-outward conformational change at TM2ab. Our structural analysis also explains the strong connection between SEA0400 binding and the Na^+^-dependent inactivation - SEA0400 is ineffective in an exchanger lacking Na^+^-dependent inactivation whereas enhancing the extent of Na^+^-dependent inactivation increases the affinity for SEA0400. As SEA0400 binding traps NCX1 in an inward-facing state that facilitates the formation of the inactivation assembly, the interaction between XIP and the TM domain in the assembly can in turn stabilize SEA0400 binding by sealing the drug-binding pocket from the cytosol. When XIP is removed as in the chymotrypsin-treated NCX1 or when the inactivation assembly cannot form as in the NCX1 at forward exchange mode, the SEA0400-binding pocket becomes exposed to the cytosol, mitigating the inhibition efficacy of SEA0400^50,51^.

While we expect the long-chain PIP_2_ to bind in the same location as PIP_2_ diC8, it is unclear how it exacerbates the destabilization effect on the inactivation assembly. The long acyl chain of native PIP_2_ may engage in some interactions with the TM module of NCX1 that are not present in the short-chain lipid, rendering a higher affinity binding and more profound destabilization effect on the inactivation assembly than PIP_2_ diC8. The acyl-chain length-dependent PIP_2_ activation is consistent with some previous studies. Before PIP_2_ was demonstrated to regulate NCX, some earlier studies showed that negatively charged long-chain lipids such as phosphatidylserine (PS) or phosphatidic acid (PA) could have the same potentiation effects on NCX1 as PIP_2_^56,57^. Furthermore, long-chain acyl-CoAs could also have the same potentiation effects on NCX as PIP_2_^58^. All these studies demonstrated that activation of NCX by the anionic lipids depends on their chain length with the short chain being ineffective or less effective. These findings have two implications. First, it is the negative surface charge rather than the specific IP_3_ head group of the lipid that is important for stimulating NCX1 activity, suggesting non-specific electrostatic interactions between the negatively charged lipids and those positively charged residues at the binding site. Second, a longer acyl chain is required for the high-affinity binding of PIP_2_ or negatively charged lipids. In the PIP_2_ diC8-bound structure, the tail of the acyl chain is positioned right at the fenestration of NCX1 that serves as the portal for the SEA0400 binding. Two pieces of evidence lead us to suggest that the tail of the long acyl chain from a native lipid can enter the same binding pocket for SEA0400 and thereby rendering its higher affinity binding than a shorter chain lipid. Firstly, a docking analysis^59,60^ of both long-chain and short-chain PIP_2_ at the binding site showed that the tail portion of the acyl chain from the native brain PIP_2_ inserts into the SEA0400-binding pocket through the fenestration of NCX1 in all docking models with highest calculated affinity (Fig. S6A). This pocket is not reachable for PIP_2_ diC8 due to its shorter chain length (Fig. S6B). Second, a stretch of density likely from a native lipid acyl chain is observed in the apo NCX1 structure inside the SEA0400-binding pocket (Fig. S6C), indicating its accessibility to a long-chain lipid. Thus, the accommodation of the long acyl chain in the open pocket of NCX1 through the fenestration in the middle of the membrane likely contributes to the high affinity binding of native lipids.

## Methods

### Protein expression and purification

The expression and purification of the HsNCX1 (cardiac isoform NCX1.1, indicated as HsNCX1 or NCX1 throughout the manuscript) were carried out as described previously^32^. Truncated HsNCX1 (Δ341-365aa) containing a C-terminal Strep-tag was cloned into a pEZT-BM vector and baculoviruses were produced in *Sf9* cells^61^. For protein expression, cultured Expi293F GnTI-cells were infected with the baculoviruses at a ratio of 1:20 (virus: GnTI-, v/v) for 10 hours. 10 mM sodium butyrate was then introduced to boost protein expression level and cells were cultured in suspension at 30°C for another 60 hours and harvested by centrifugation at 4,000 g for 15 mins. All purification procedures were carried out at 4°C. The cell pellet was resuspended in lysis buffer (25 mM HEPES pH 7.4, 300 mM NaCl, 2 μg/ml DNase I, 0.5 μg/ml pepstatin, 2 μg/ml leupeptin, 1 μg/ml aprotinin, and 0.1 mM PMSF) and homogenized by sonication. NCX1 was extracted with 2% (w/v) N-dodecyl-β-D-maltopyranoside (DDM, Anatrace) supplemented with 0.2% (w/v) cholesteryl hemisuccinate (CHS, Sigma Aldrich) by gentle agitation for 2 hours and supernatant collected by centrifugation at 40,000 g for 30 mins was incubated with Strep-Tactin affinity resin (IBA) for 1 hour. The resin was then collected on a disposable gravity column (Bio-Rad) and washed with 30 column volumes of buffer A (25 mM HEPES pH 7.4, 200 mM NaCl, 0.06% digitonin). NCX1 was eluted in buffer A supplemented with 50 mM biotin and further purified by size-exclusion chromatography on a Superdex200 10/300 GL column (GE Healthcare). For the generation of NCX1-Fab 2E4 complex, NCX1 was incubated with purified Fab in a molar ratio of 1:1.2 (NCX1: Fab 2E4) for 2 hours and further purified by size-exclusion chromatography in buffer A. The peak fractions were collected and concentrated to ∼5-6 mg/ml for cryo-EM analysis. To prepare the protein samples in complex with various compounds, 0.9 mM SEA0400, 0.47 mM PI(4,5)P_2_ diC8 or 0.42 mM brain PI(4,5)P_2_ were added to the protein samples 2 hours before grid preparation.

Expi293F GnTI-cells were purchased from and authenticated by Thermo Fisher Scientific. The cell lines were tested negative for mycoplasma contamination.

### Cryo-EM sample preparation and data acquisition

HsNCX1-Fab 2E4 samples (∼5-6 mg/ml) in various conditions were applied to a glow-discharged Quantifoil R1.2/1.3 300-mesh gold holey carbon grid (Quantifoil, Micro Tools GmbH, Germany), blotted for 4.0 s under 100% humidity at 4 °C and plunged into liquid ethane using a Mark IV Vitrobot (FEI). For the SEA0400-bound NCX1-Fab 2E4, raw movies were acquired on a Titan Krios microscope (FEI) operated at 300 kV with a K3 camera (Gatan) at 0.83 Å per pixel and a nominal defocus range of −0.9 to −2.2 μm. Each movie was recorded for about 5 s in 60 subframes with a total dose of 60 e^-^/Å^2^. For other samples, raw movies were acquired on a Titan Krios microscope operated at 300 kV with a Falcon 4i (Thermo Fisher Scientific) at 0.738 Å per pixel and a nominal defocus range of −0.8 to −1.8 μm. Each movie was recorded for 4 s with a total dose of 60 e^-^/Å^2^.

### Image processing

Cryo-EM data were processed following the general scheme described below with some modifications to different datasets (Figs. S1, S2, and S4). First, movie frames were motion-corrected and dose-weighted using MotionCor2^62^. The CTF parameters of the micrographs were estimated using the GCTF program^63^. After CTF estimation, micrographs were manually inspected to remove images with bad defocus values and ice contamination. Particles were picked using program Gautomatch (Kai Zhang, https://www.mrc-lmb.cam.ac.uk/kzhang/) or crYOLO^64^ and extracted with a binning factor of 3 in RELION^65,66^. Extracted particles were subjected to 2D classification, ab initio modeling, and 3D classification. The particles from the best-resolving 3D class were then re-extracted with the original pixel size and subjected to heterogenous 3D refinement, non-uniform refinement, CTF refinement, and local refinement in cryoSPARC^67^. The quality of the EM density maps for the transmembrane (TM) and cytosolic domains was further improved through focused refinement, allowing for accurate model building for a major part of the protein. For the dataset of NCX1 in complex with brain PI(4,5)P_2_, the maps of apo inactive NCX1 (PDB 8SGJ) and Ca²⁺-bound active NCX1 (PDB 8SGT) are used as references for heterogeneous refinement. Due to the highly dynamic nature of the protein samples, the particles sorted into the active state produce a map with very low resolution. All resolution was reported according to the gold-standard Fourier shell correlation (FSC) using the 0.143 criterion^68^. Local resolution was estimated using cryoSPARC.

### Model building, refinement, and validation

The EM maps of HsNCX1 in the SEA0400-bound and PI(4,5)P_2_ diC8-bound states show high-quality density and model building is facilitated by previous apo NCX1 structure (PDB 8SGJ)^32^. Models were manually adjusted in Coot ^69^ and refined against maps using the phenix.real_space_refine with secondary structure restraints applied^70^. The final NCX1structural model contains residues 17-248, 370-467, 482-644, 652-698, 707-718, and 738-935. The EM map of HsNCX1 in complex with brain PI(4,5)P_2_ is relatively poor. The Ca^2+^-bound activated NCX1 structure (PDB 8SGT) is directly docked into the EM map without adjustment.

The statistics of the geometries of the models were generated using MolProbity^71^. All the figures were prepared in PyMol (Schrödinger, LLC.), Chimera^72^, and ChimeraX^73^.

### Electrophysiological experiments

The wild-type HsNCX1 and its mutants were cloned into a pGEMHE vector and expressed in oocytes for electrophysiological recordings. RNA was synthesized using mMessage mMachine (Ambion) and injected into Xenopus laevis oocytes as described in^74^. Oocytes were isolated from at least 3 different frogs and kept at 18°C for 4-7 days. Outward HsNCX1 currents were recorded using the giant patch technique in the inside-out configuration. Each data point shown in this study represents a recording obtained from a single oocyte. The external solution (pipette solution) contained the following (mM): 100 CsOH (cesium hydroxide), 10 HEPES (4-(2-hydroxyethyl)-1-piperazineethanesulfonic acid), 20 TEAOH (tetraethyl-ammonium hydroxide), 0.2 niflumic acid, 0.2 ouabain, 8 Ca(OH)_2_ (calcium hydroxide), pH = 7 (using MES, (2-(*N*-morpholino) ethanesulfonic acid)); bath solution (mM): 100 CsOH or 100 NaOH (sodium hydroxide), 20 TEAOH, 10 HEPES, 10 EGTA(ethylene glycol-bis(β-aminoethyl ether)-*N*,*N*,*N*’,*N*’-tetra acetic acid) or HEDTA (N-(2-Hydroxyethyl) ethylenediamine-N,N’,N’-triacetic acid) and different Ca(OH)_2_ concentrations to obtain the desired final free Ca^2+^ concentrations, pH = 7 (using MES). Free Ca^2+^ concentrations were calculated according to WEBMAXc program and confirmed with a Ca^2+^ electrode.

HsNCX1 currents were evoked by the rapid replacement of 100 mM Cs^+^ with 100 mM Na^+^, using a computer-controlled 20-channel solution switcher. As HsNCX1 does not transport Cs^+^, there is no current and only upon application of Na^+^ does the exchange cycle initiate. Data were acquired at 4 ms/point and filtered at 50 Hz using an 8-pole Bessel filter. Experiments were performed at 35°C and at a holding potential of 0 mV. The effects of the Na^+^-dependent inactivation were analyzed by measuring fractional currents calculated as the ratio of the steady-state current to the peak current (fractional activity). All *P* values were calculated using an unpaired, two-sided Welch’s *t*-test.

PI(4,5)P_2_ diC8 (Phosphatidylinositol 4,5-bisphosphate diC8, Echelon Bioscience) and brain PI(4,5)P_2_ (L-α-phosphatidylinositol-4,5-bisphosphate, Brain, Porcine, Avanti Polar Lipids) were dissolved in water and kept as stock at −20 °C. Immediately prior to recordings, PI(4,5)P_2_ was diluted in the bath solution to 10 µM final concentration and perfused cytosolically for the indicated time.

## Data availability

The cryo-EM density maps of the human NCX1 have been deposited in the Electron Microscopy Data Bank under accession numbers EMD-40456 for the SEA0400-bound state and EMD-60921 for the PI(4,5)P_2_ diC8-bound state, respectively. Atomic coordinates have been deposited in the Protein Data Bank under accession numbers 8SGI for SEA0400-bound structure and 9IV8 for the PI(4,5)P_2_ diC8-bound structure.

## Acknowledgments

Single particle cryo-EM data were collected at the University of Texas Southwestern Medical Center Cryo-EM Facility which is funded by the CPRIT Core Facility Support Award RP170644. Cryo-EM sample grids were prepared at the Structural Biology Laboratory at UT Southwestern Medical Center which is partially supported by grant RP170644 from CPRIT. This work was supported in part by the Howard Hughes Medical Institute (to Y.J.) and by grants from the National Institute of Health (R35GM140892 to Y.J. and R01HL152296 to M.O.), and the Welch Foundation (Grant I-1578 to Y.J.).

## Author contributions

J.X. prepared the samples and performed data acquisition, image processing, and structure determination; M.O., S.J., N.A., and W.Z. performed electrophysiology recording; Y.J. supervised the work; all authors participated in research design, data analysis, discussion, and manuscript preparation.

---

**Fig. S1.**
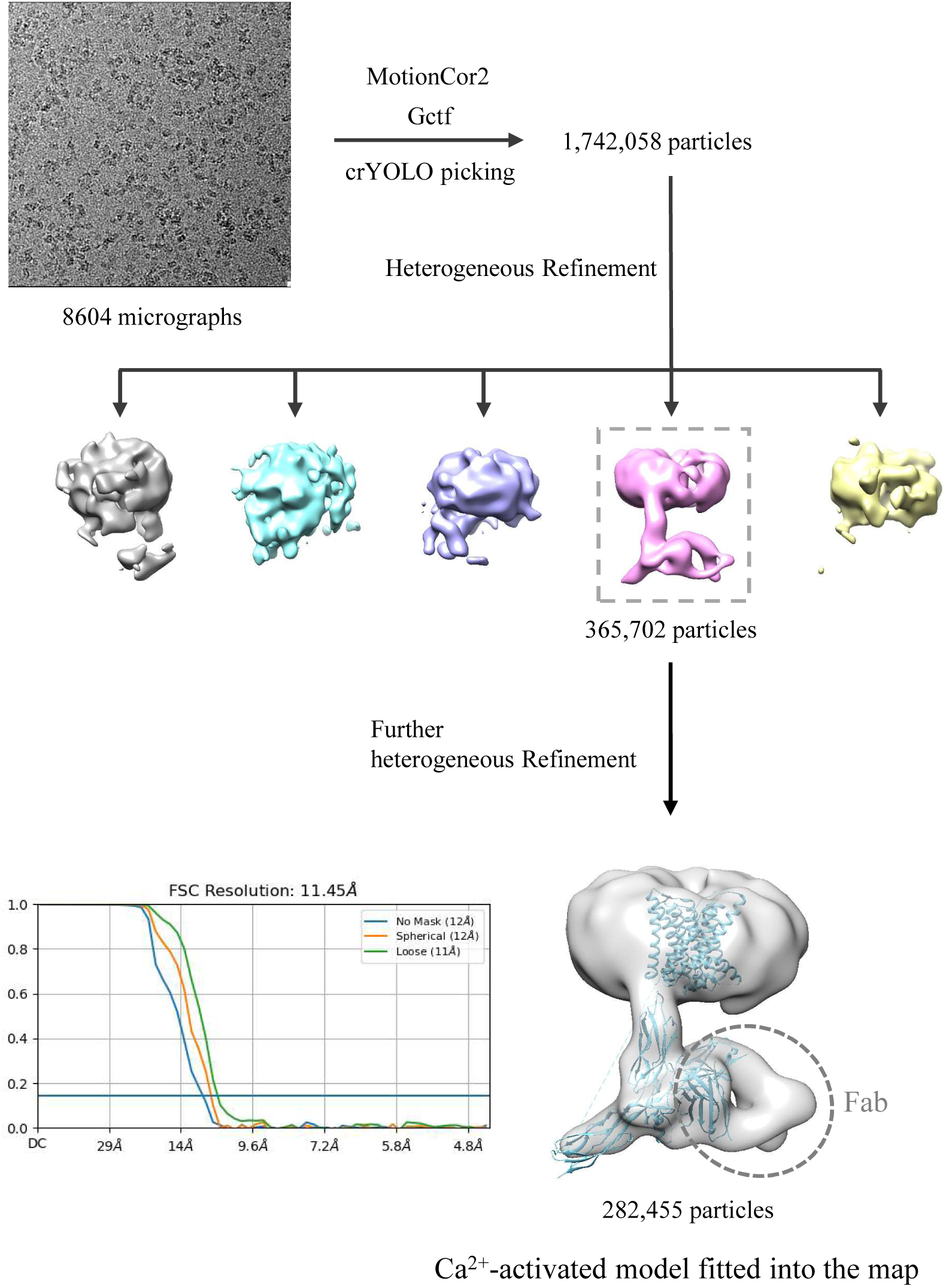
Cryo-EM data processing of HsNCX1 in the presence of long-chain porcine brain PIP_2_. The structural model of Ca2+-activated HsNCX1 from a previous study (PDB 8SGT) was directly fitted into the low-resolution EM-map (∼11.5 Å). The Fab fragment from a monoclonal antibody against NCX1 was used as a fiducial marker to facilitate the single-particle alignment.

**Fig. S2.**
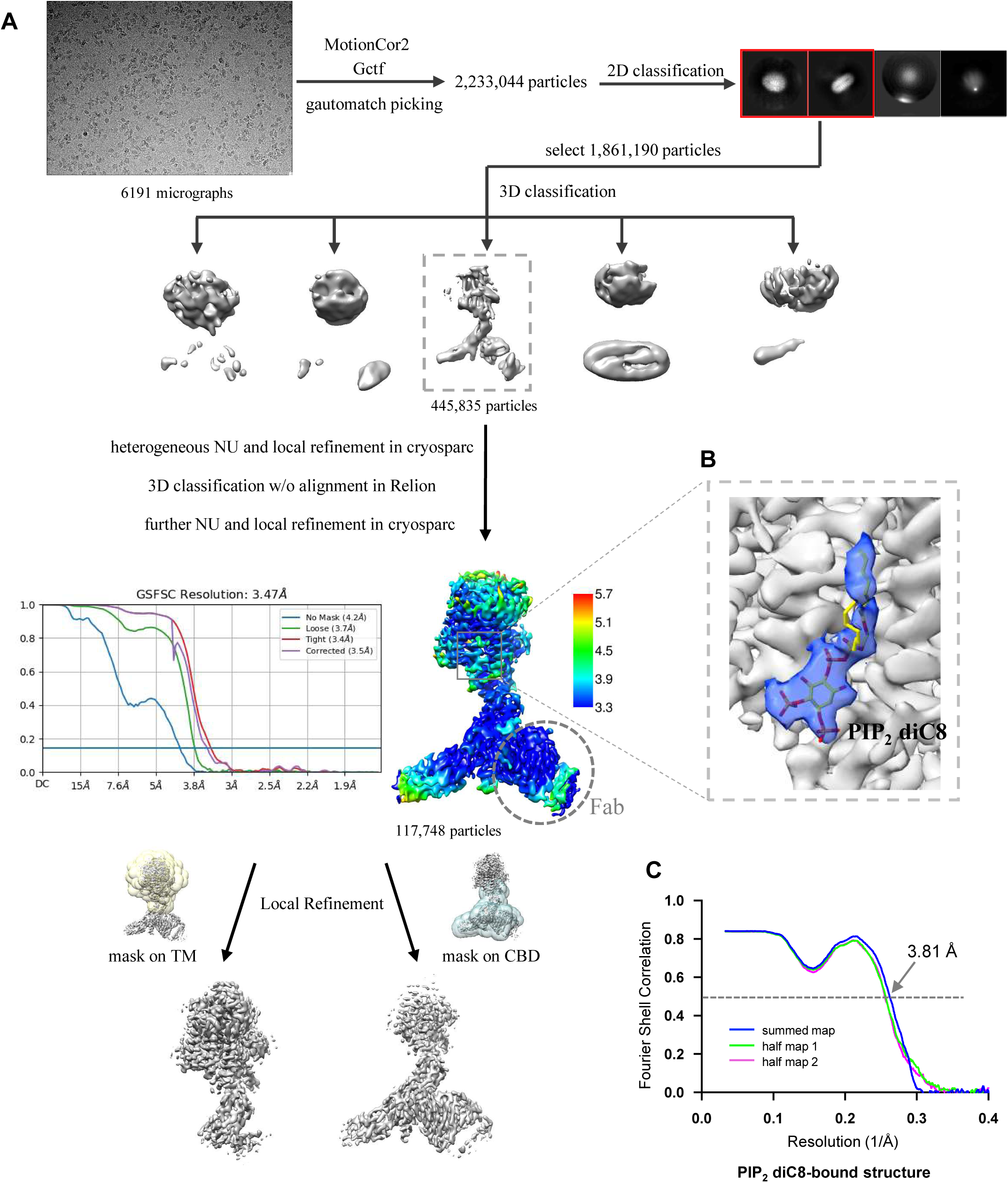
Structure determination of HsNCX1-PIP_2_ diC8 complex. **(A)** Cryo-EM data processing of HsNCX1 in complex with short-chain PIP_2_ diC8. The Fab fragment from a monoclonal antibody against NCX1 was used as a fiducial marker to facilitate the single-particle alignment. **(B)** Zoomed-in view of density map of the bound PIP_2_ diC8 contoured at the threshold level of 0.35 using ChimeraX. **(C)** The Fourier shell correlation (FSC) curves for cross-validation between the maps and the models.

**Fig. S3.**
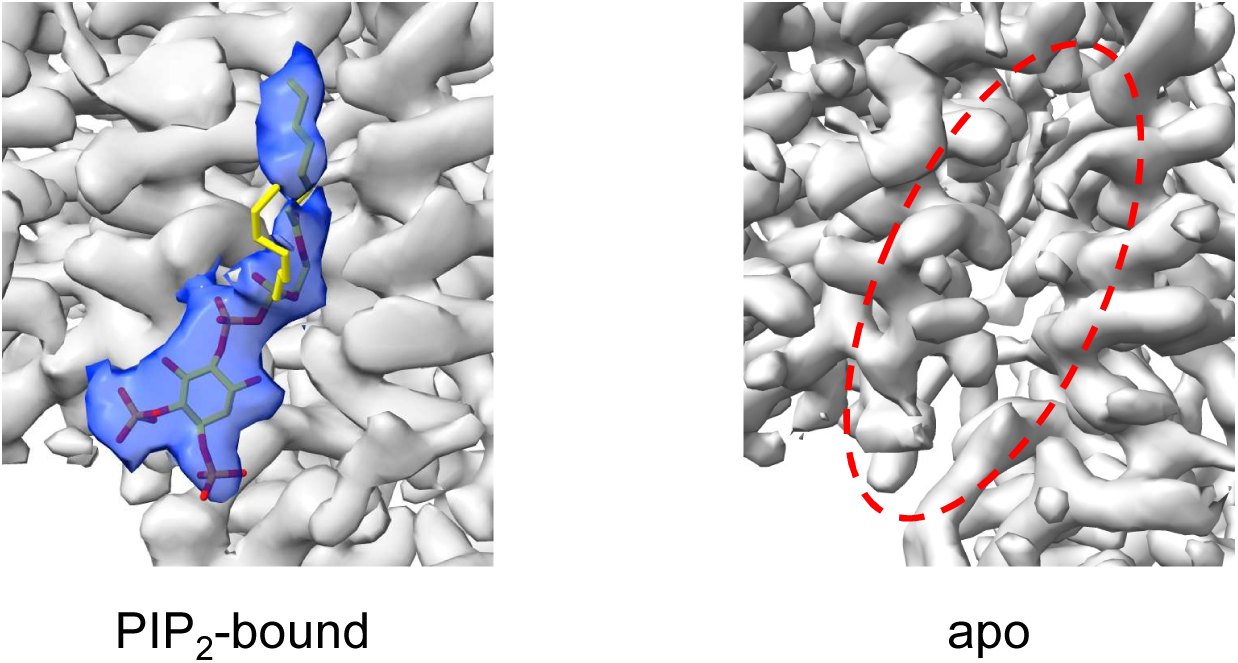
Side-by-side comparison of the densities at the PIP_2_-binding site between the PIP_2_-bound structure (EMD-60921) and the apo structure (EMD-40457). The local-refined maps focused on the TM domain are used in the comparison. The density map of the bound PIP_2_ is contoured at a threshold level of 0.35 using ChimeraX.

**Fig. S4.**
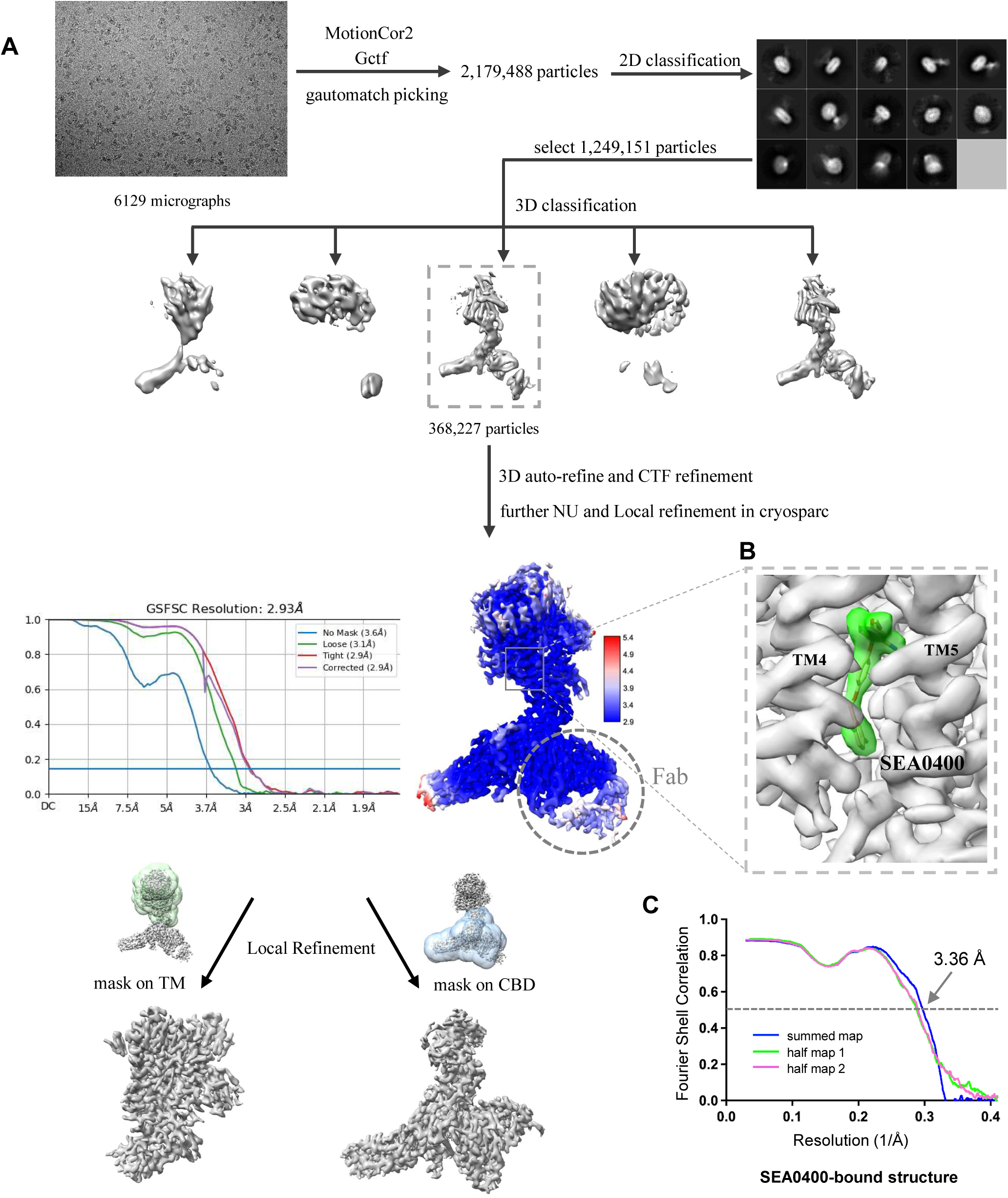
Structure determination of HsNCX1-SEA0400 complex. **(A)** Cryo-EM data processing scheme of HsNCX1 in complex with SEA0400 inhibitor. The Fab fragment from a monoclonal antibody against NCX1 was used as a fiducial marker to facilitate the single-particle alignment. **(B)** Zoomed-in view of density map of the bound SEA0400 inhibitor contoured at the threshold level of 0.52 using ChimeraX. **(C)** The Fourier shell correlation (FSC) curves for cross-validation between the maps and the models.

**Fig. S5.**
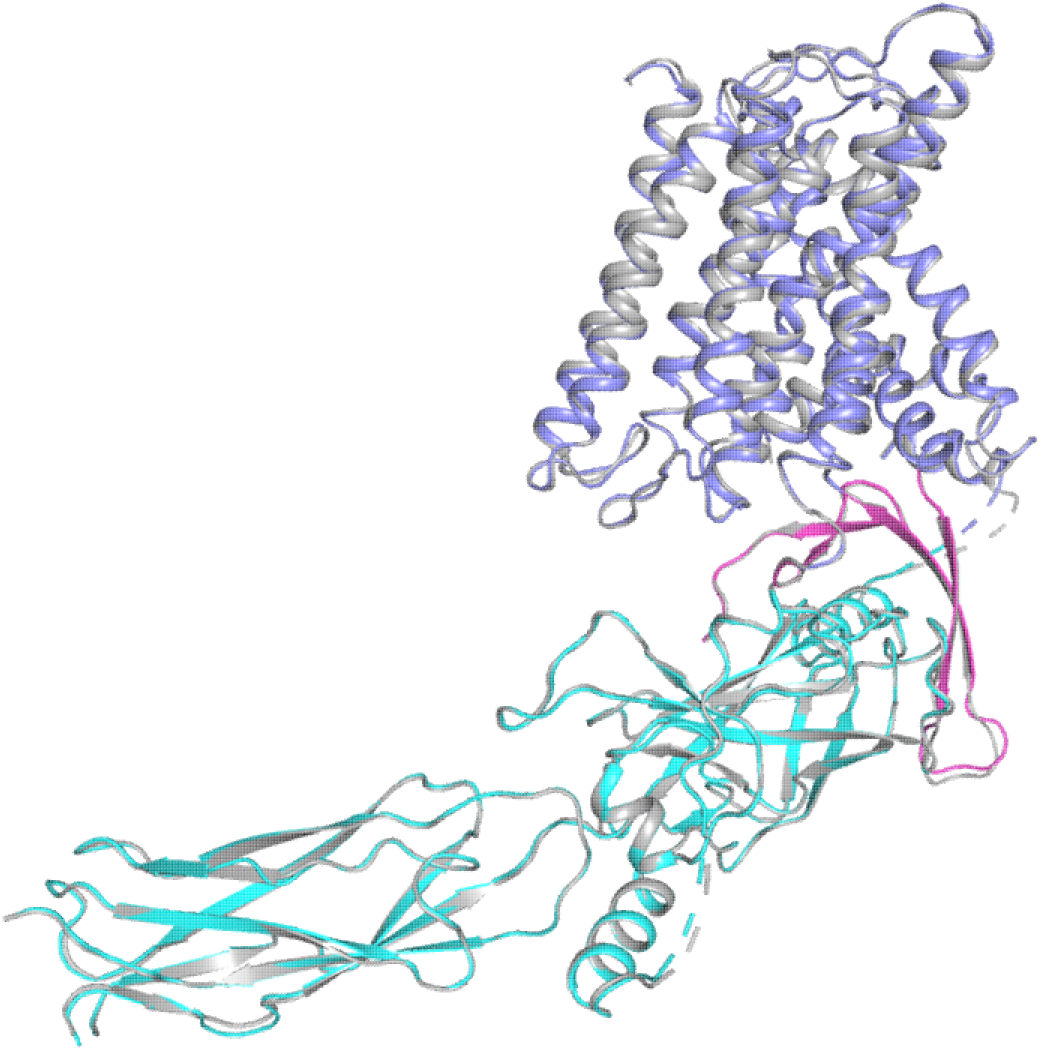
Structural comparison between the apo (grey) and SEA0400-bound (color) HsNCX1.

**Fig. S6.**
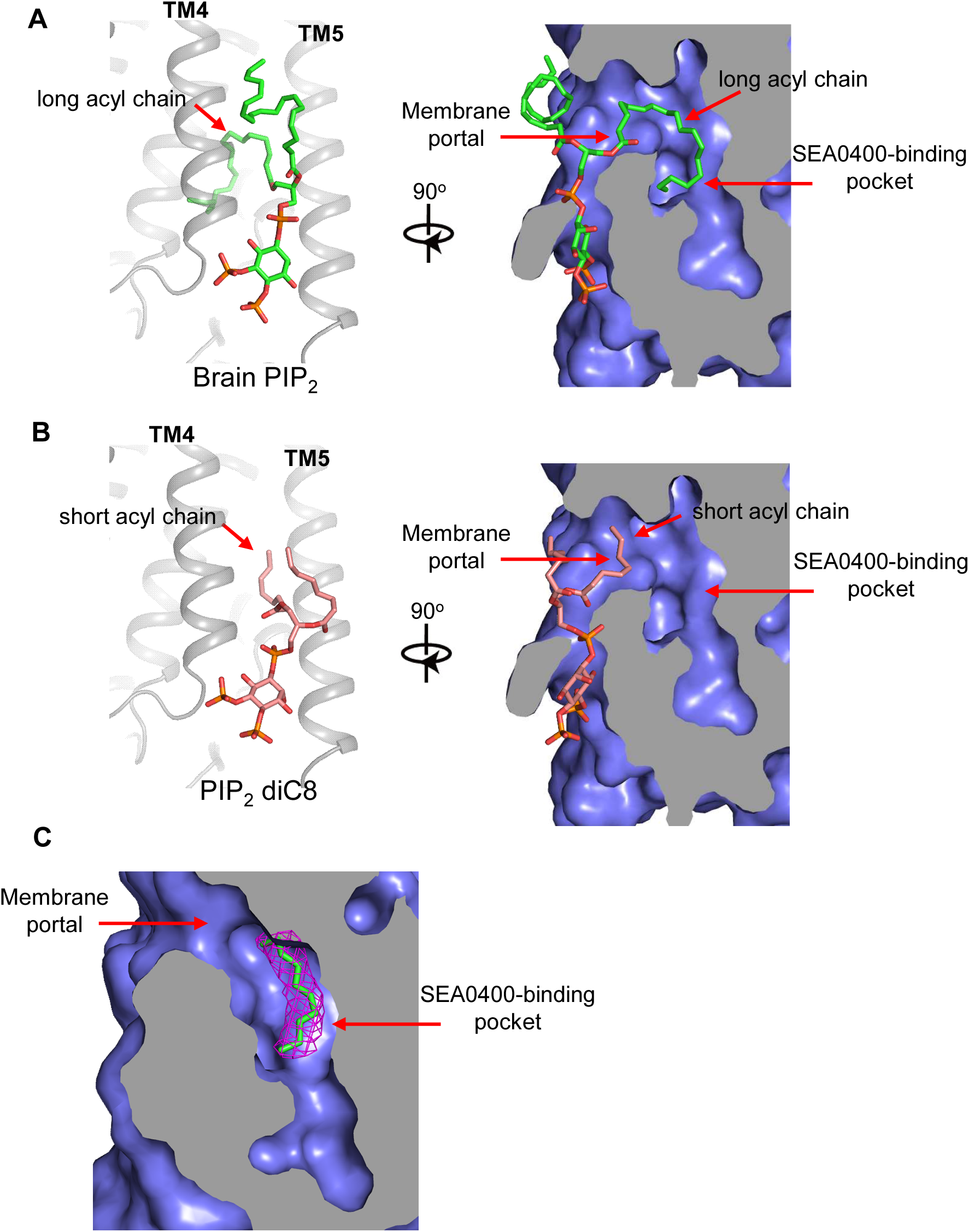
Proposed structural basis underlying the different binding affinity between long and short-chain PIP_2_. (A) A docking model for native brain PIP_2_ binding in NCX1 showing the insertion of its long acyl chain into the SEA0400-binding pocket. (B) A docking model for short-chain PIP_2_ diC8 binding in NCX1 showing that the SEA0400-binding pocket is not accessible to shorter acyl chain. (C) The density (red mesh, contoured at 6*σ*) of an acyl chain likely from a native lipid is observed in the SEA0400-binding pocket of the apo NCX1 structure (EMD-40457, local-refined map at the TM domain)

**Table S1.** Cryo-EM data collection and model statistics.

|  |  |  |
| --- | --- | --- |
| **Sample preparation conditions** | 25 mM Hepes pH 7.4,<br>200 mM NaCl<br>0.9 mM SEA0400 | 25 mM Hepes pH 7.4,<br>200 mM NaCl,<br>0.47 mM PI(4,5)P<sub>2</sub> diC8 |
|  | **SEA0400-bound state**<br>(EMD-40456,<br>PDB 8SGI) | **PI(4,5)P<sub>2</sub> diC8-bound state**<br>(EMD-60921,<br>PDB 9IV8) |
| **Data collection and processing** |  |  |
| Magnification | 105k | 105k |
| Voltage (kV) | 300 | 300 |
| Electron exposure (e<sup>-</sup>/Å<sup>2</sup>) | 60 | 60 |
| Defocus range (μm) | -0.9 - -2.2 | -0.9 - -2.2 |
| Pixel size (Å) | 0.83 | 0.84 |
| Symmetry imposed | C1 | C1 |
| Initial particle images (no.) | 1,249,151 | 2,233,044 |
| Final particle images (no.) | 368,227 | 117,748 |
| Map resolution (Å) | 2.93 | 3.47 |
| FSC threshold | 0.143 | 0.143 |
| **Refinement** |  |  |
| Initial model used<br>(PDB code) | 8SGJ | 8SGJ |
| Model resolution (Å) | 3.36 | 3.81 |
| FSC threshold | 0.5 | 0.5 |
| Model composition |  |  |
| Non-hydrogen atoms | 7667 | 5947 |
| Protein residues | 982 | 750 |
| Ligands | 3: Na<br>6: Ca<br>1: H<sub>2</sub>O<br>1: SEA0400 | 5: Ca<br>1: PI(4,5)P<sub>2</sub> diC8 |
| B factors (Å<sup>2</sup>) |  |  |
| Protein | 66.21 | 50.47 |
| Ligands | 58.55 | 106.20 |
| R.m.s. deviations |  |  |
| Bond lengths (Å) | 0.005 | 0.004 |
| Bond angles (°) | 0.696 | 0.679 |
| Validation |  |  |
| MolProbity score | 1.36 | 1.28 |
| Clashscore | 5.24 | 5.22 |
| Poor rotamers (%) | 0 | 0 |
| Ramachandran plot |  |  |
| Favored (%) | 97.62 | 98.24 |
| Allowed (%) | 2.38 | 1.76 |
| Disallowed (%) | 0 | 0 |

